# Selective vulnerability and resilience to Alzheimer’s disease tauopathy as a function of genes and the connectome

**DOI:** 10.1101/2024.03.04.583403

**Authors:** Chaitali Anand, Justin Torok, Farras Abdelnour, Pedro D. Maia, Ashish Raj

## Abstract

Brain regions in Alzheimer’s (AD) exhibit distinct vulnerability to the disease’s hallmark pathology, with the entorhinal cortex and hippocampus succumbing early to tau tangles while others like primary sensory cortices remain resilient. The quest to understand how local/regional genetic factors, pathogenesis, and network-mediated spread of pathology together govern this selective vulnerability (SV) or resilience (SR) is ongoing. Although many risk genes in AD are known from gene association and transgenic studies, it is still not known whether and how their baseline expression signatures confer SV or SR to brain structures. Prior analyses have yielded conflicting results, pointing to a disconnect between the location of genetic risk factors and downstream tau pathology. We hypothesize that a full accounting of genes’ role in mediating SV/SR would require the modeling of network-based vulnerability, whereby tau misfolds, aggregates, and propagates along fiber projections.

We therefore employed an extended network diffusion model (eNDM) and tested it on tau pathology PET data from 196 AD patients from the Alzheimer’s Disease Neuroimaging Initiative (ADNI). Thus the fitted eNDM model becomes a reference process from which to assess the role of innate genetic factors. Using the residual (observed *–* model-predicted) tau as a novel target outcome, we obtained its association with 100 top AD risk-genes, whose baseline spatial transcriptional profiles were obtained from the Allen Human Brain Atlas (AHBA). We found that while many risk genes at baseline showed a strong association with regional tau, many more showed a stronger association with residual tau. This suggests that both direct vulnerability, related to the network, as well as network-independent vulnerability, are conferred by risk genes. We then classified risk genes into four classes: network-related SV (SV-NR), network-independent SV (SV-NI), network-related SR (SR-NR), and network-independent SR (SR-NI). Each class has a distinct spatial signature and associated vulnerability to tau. Remarkably, we found from gene-ontology analyses, that genes in these classes were enriched in distinct functional processes and encompassed different functional networks. These findings offer new insights into the factors governing innate vulnerability or resilience in AD pathophysiology and may prove helpful in identifying potential intervention targets.

## 1 Introduction

Extracellular amyloid-β (Aβ) plaques and intracellular tau neurofibrillary tangles, the pathological hallmarks of Alzheimer’s disease (AD), follow a stereotyped pattern of progression to distinct brain regions characterized by varying levels of vulnerability to AD^1^. Selective vulnerability (SV) is a norm rather than an exception in progressive neurodegeneration^2^, where certain brain regions are more susceptible to damage and degeneration than others. SV may be due to various factors such as the distribution of specific receptors, cellular vulnerability to oxidative stress, and the regional expression of genes associated with pathology^3, 4^. In AD, SV is more closely linked to tau pathology^3^, since tau accumulates in discrete brain regions and coincides with regions that exhibit marked neurodegeneration, whereas Aβ plaques are relatively widespread throughout the neocortex. Neurons vulnerable to the accumulation of pathological tau and lost early in the disease include large pyramidal neurons in layer II of the entorhinal cortex (EC), the subiculum, and the hippocampal CA1 region; cholinergic neurons in the basal forebrain; and noradrenergic neurons in the locus coeruleus. On the other hand, granule neurons in the dentate gyrus, deeper layer III, and layers V and VI of the EC as well as cortical interneurons are spared in early AD^3^. There is thus a time-dependent component to SV, where the EC, hippocampus, and neocortical association areas are affected early, with the primary sensory cortex being resistant to degeneration until later^5^, these may be called “selectively resilient (SR)” to pathological insults. Note that ‘resilience to AD’ here does not indicate cognitive resilience to the disease, but rather ‘pathological resilience’, which points to the capacity of certain brain regions to halt or delay the accumulation of AD pathology.

Although the etiology of SV and SR is not clearly known, current hypotheses have focused on two general concepts and causative mechanisms: (a) cell-autonomous factors (cell type, cytoarchitectural, genetic or molecular factors)^6^ and (b) non-cell autonomous factors (processes dependent on cell-cell communication, network connectivity, and topology)^7^. Cell-autonomous factors represent the most direct and plausible mechanisms underlying regional SV or SR. The variability in the regional levels of pathogenic proteins is largely governed by the genetic environment of that region^8^. AD is a clinically heterogeneous neurodegenerative disease with a strong genetic component. Over 40 risk loci for late-onset AD have been identified via large genome-wide association studies (GWAS), most of which are common variants with small effect sizes^9^. It is very likely that certain genes confer SV or SR causing some regions to be more vulnerable to pathology even before disease onset. From a biological perspective, there are several ways in which *baseline* healthy gene expression may play a role in directing the future occurrence and spread of pathogenic species. A high baseline expression of risk genes in a given region may effectively eventually increase the local accumulation rate of pathology (tau, for instance) rendering the region selectively vulnerable. Conversely, some genes can contribute to SR and have a protective role instead, such as through the enhancement of clearance at a regional level resulting in regions with less pathological burden. The genetic/molecular explanation of SV/SR would necessitate a direct association between genes and AD topography. Remarkably, however, the spatial distribution of vulnerable regions bears little relation to that of the associated genetic/molecular factors^10–13^; this notable dissociation between upstream genes and downstream pathology has been called one of the key mysteries of neurodegenerative diseases^13, 14^.

In this study, we put forth the hypothesis that this dissociation may be caused, and hence explained, by the prominent underlying role of non-cell-autonomous mechanisms, specifically that of trans-neuronal transmission of tau pathology. Indeed, tau is increasingly thought to spread along the brain’s anatomic connectivity network^15–17^. Injection of synthetic tau fibrils in the mouse brain induces further misfolding and propagation to distant but connected regions^18, 19^, implicating non-cell-autonomous, network connectivity-mediated factors in pathology spread. The spatiotemporal dynamics of tau spread have been modeled as a diffusive process between connected brain regions by us and others via a system of differential equations called the Network Diffusion Model (NDM)^10, 17, 20^ and its extension called Nexopathy in silico or “Nexis” model in Anand *et al*^21^. In these models pathology progression is primarily mediated by the brain’s anatomic network - a concept we call “network vulnerability or resilience”. Due to the underlying network-mediated spread and aggregation of tau pathology, exploration of cell-autonomous genetic/molecular factors of SV/SR requires a more careful and joint consideration of both network effects and the role of genes and cells - a concept popularized by the so-called “molecular nexopathy” paradigm^21, 22^.

We hypothesize that this joint analysis will reveal the role of genes unambiguously and in relation to the dominant mediating role of the network. We wish not only to identify candidate genes that contribute to SV/SR in this joint analysis but also classify them according to their exact relationship with network vulnerability or resilience. Therefore we obtained a mathematical “extended network diffusion model” (eNDM) of network vulnerability derived from a simplified and humanized version of the Nexis model previously successful in predicting tau progression in mesoscale mouse studies^21^. We fitted the eNDM to individual Alzheimer patients’ tau PET data and achieved excellent match both qualitatively and quantitatively. Thus the fitted eNDM model becomes a reference process from which to assess the role of innate genetic factors. An overview of our data processing and analysis pipeline is shown in (Figure 1).

**Figure 1:**
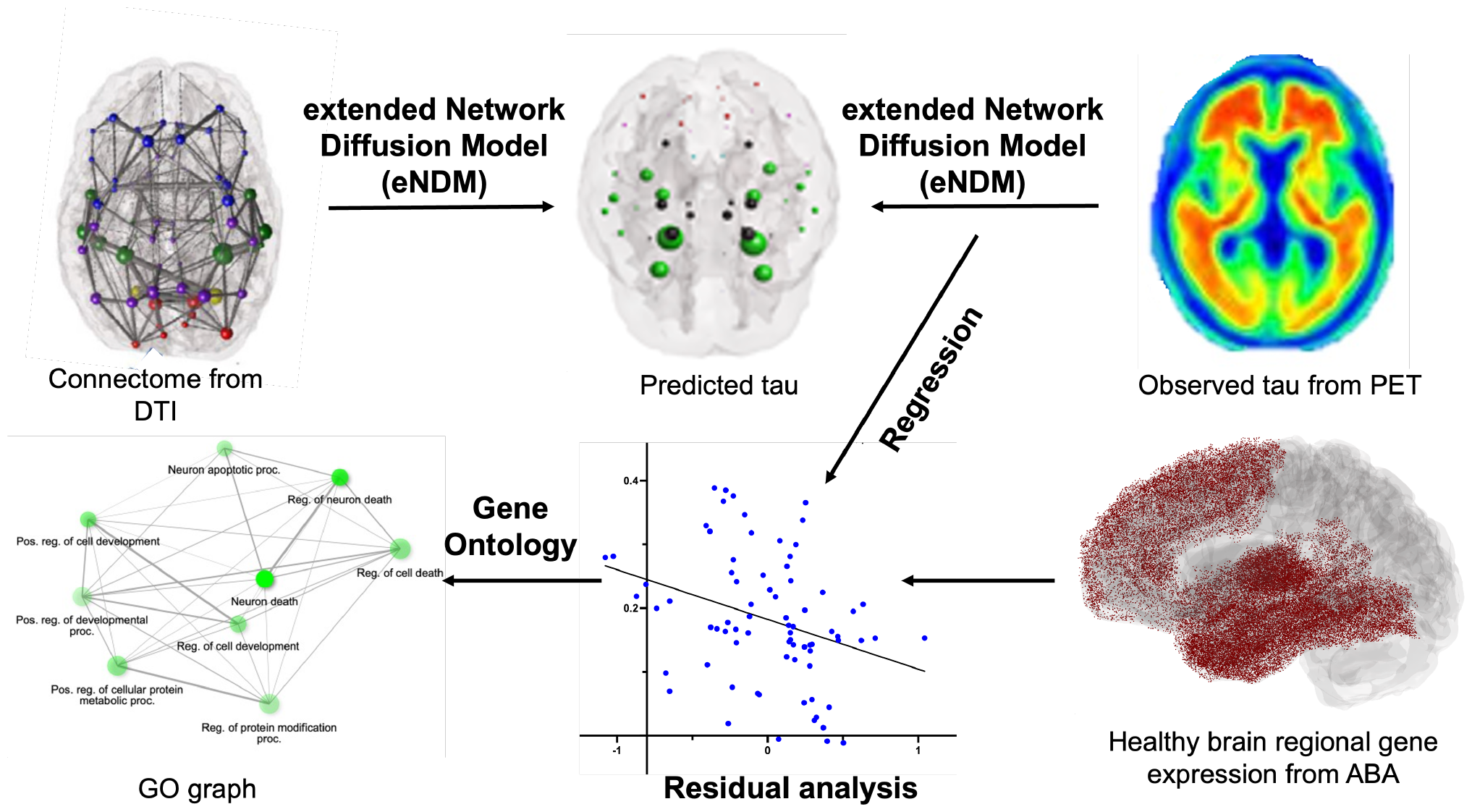
Workflow of selective vulnerability and resilience analysis. Structural tractography data from DTI analysis and regional tau PET MRI data (observed tau) are entered into the extended Network Diffusion Model (eNDM) to calculate predicted tau. Linear regression analysis between the observed (PET MRI) tau and eNDM-predicted tau provides residual tau levels. Baseline regional gene expression data obtained from the Allen Human Brain Atlas (AHBA) are correlated with regional observed and residual tau levels to identify genes in the selective vulnerability and selective resilience (SV and SR) groups. This is followed by gene ontology analysis (GO) on the gene sets identified above.

We found that most known risk genes act to enhance SV or SR in two modes: either dependent or independent of network transmission. This classification of risk genes into 4 classes adds new avenues for assessing their impact on AD pathophysiology. Many genes that are strong risk factors show no association with regional tau directly, but a strong association once the network vulnerability has been accounted for. Remarkably, we found, via detailed gene ontology, that the identified gene classes have distinct and sometimes surprising functional enrichment patterns. Note that throughout this manuscript, SV and SR refer to *innate* (baseline gene expression-mediated) selective vulnerability and resilience, as opposed to disease-associated changes in gene expression. Such expression patterns may lay down vulnerability networks for AD pathogenesis to occur once triggered by appropriate initiating factors^23^. They can also point to factors that might contribute to pathology resilience in certain other networks. Understanding the underlying factors that govern innate regional vulnerability is not only an important scientific goal but will also aid in identifying targets for intervention, an effort that has gained urgency with the rising prevalence of AD and few therapeutic options for slowing its progression.

## 2 Materials and Methods

The goal of this study was to establish a relation between regional baseline expression of AD-related genes and regional tau levels. This relationship was also extended to regional network diffusion model-predicted and residual tau levels.

### 2.1 Data

#### Patients

Data used in this study were obtained from the Alzheimer’s Disease Neuroimaging Initiative (ADNI3) database (adni.loni.usc.edu). Empirical AV1451-PET (“tau-PET”) imaging data were obtained from a sample of 196 patients consisting of patients with AD and those in the early and late stages of mild cognitive impairment (EMCI and LMCI). The final sample was composed of 102 patients with EMCI (64 men/38 women, mean age 77.73 *±* 7.17), 47 patients with LMCI (29 men/18 women, mean age 75.94 *±* 7.56), and 47 patients with AD (25 men/22 women, mean age 78.43 *±* 9.17).

#### Brain regional parcellation and tau co-registration

High-resolution T1-weighted sagittal MR images of patients’ brains were obtained from the ADNI website (https://adni.loni.usc.edu/data-samples/access-data/). These images were acquired using the 3D MPRAGE sequence with specific parameters: an 8-channel coil, TR (Repetition Time) of 400 ms, minimum full TE (Echo Time), 11 degrees flip angle, slice thickness of 1.2 mm, resolution of 256 × 256 mm, and a FOV (Field of View) of 26 cm. Briefly, the ADNI image processing pipeline was as follows. The T1 images underwent normalization to the Montreal Neurological Institute (MNI) space and segmentation using SPM8’s unified coregistration and segmentation scheme (https://www.fil.ion.ucl.ac.uk/spm/). Grey matter (GM) was divided into 86 regions of interest (ROIs) based on the Desikan-Killiany (DK) atlas^24^. Post-processed tau-PET images were retrieved from ADNI and underwent a series of further processing steps. Initially, these images were normalized to a common space, adjusted using cerebellar values, and resliced to ensure uniform voxel resolution. To establish a reference point, PET images from healthy controls were utilized to generate an average image, which was then normalized to the MNI space using SPM8’s linear and non-linear transformation techniques. This transformation was consistently applied to all individual PET images, aligning them with the same resolution as the 86-region DK atlas. Finally, mean AV1451 values were computed for each of the GM ROIs to quantify tau-PET signals within specific brain regions. Tau PET data are available for download from the ADNI website.

The PET data were then transformed into regional group statistics. Groupwise, unpaired t-tests were performed between the EMCI, LMCI, and AD groups for the AV1451-PET signal averages in each GM region. T-statistics were calculated per the convention, where a higher positive value signifies a greater degree of pathology. Subsequently, we scaled the t-statistic to have a mean of 1 and replaced all negative entries with 0.

#### Diffusion MRI (dMRI) and tractography processing

All processing was carried out within a custom pipeline based on the NiPype framework^25^. T1 images were segmented into grey (GM) and white matter (WM) and CSF tissue maps using SPM where T1 images were registered and transformed to MNI space. dMRI volumes were corrected for Eddy currents and small head movements by registration of the diffusion-weighted volumes to the first non-diffusion-weighted volume via an affine transformation using FSL FLIRT^26^. Skull-stripping was done using FSL BET. Details regarding this processing can be found in a previous publication^27^.

#### Structural connectivity network

Different structural connectivity networks were constructed using the same DK parcellations as above. First, we obtained publicly available dMRI data from the MGH-USC Human Connectome Project (HCP) to create an average template connectome. The HCP contains data from 418 healthy brains^28^. Subject-specific structural connectivity was computed on dMRI data as described in Abdelnour *et al* (2014) and Owen *et al* (2013)^29, 30^. *Bedpostx* was used to determine the orientation of brain fibers in conjunction with FSL FLIRT^26^. Tractography was performed using *probtrackx2* to determine the elements of the adjacency matrix. 4,000 streamlines were initiated from each seed voxel corresponding to a cortical or subcortical GM region and the number of streamlines reaching a target GM region was recorded. The weighted connection between the two structures, *c*(*i, j*), was defined as the number of streamlines initiated by voxels in region *i* that reached any voxel in region *j*, normalized by the sum of the source and target region volumes. The average connection strengths between both directions, (*c*(*i, j*) and *c*(*j, i*)), formed the undirected edges. To determine the geographic location of an edge, the top 95% nonzero voxels were computed for both edge directions and the consensus edge was defined as the union between both post-threshold sets. Details regarding this processing can be found in Verma et al^31^.

#### Regional gene expression analysis

Prominent AD genes were identified from the list of AD loci and genes compiled by Alzheimer’s Disease Sequencing Project (ADSP) Gene Verification Committee as well as referenced from various studies^32–36^. A total of 100 genes were shortlisted (**Supplementary Tables S1-4**). For each gene, healthy baseline expression levels were obtained from six postmortem brains provided by the Allen Human Brain Atlas (AHBA)^37^. Only two of the six brains included samples from the right hemisphere. Thus, analyses were conducted from microarray gene expressions obtained from these two donors with full spatial coverage.

We used a Python-based toolbox, `abagen` (version 0.1.3)^38^, to reliably and robustly process and map gene expression to the 86 regions in the DK volumetric atlas in MNI space, following previous work^39^. The `abagen` process has been described in detail elsewhere^38, 40^. Briefly, microarray probes were reannotated using data provided by Arnatkeviciute *et al*^41^. Filtering the probes based on their expression intensity relative to background noise as in Pandya *et al*^40^ yielded 30534 probes. Additional details on methods for gene expression data processing and mapping can be found in Pandya *et al*^40^. From the final regional gene expression matrix, we chose 100 AD genes giving us a gene expression table, *G* (86 × 100) (**Supplementary Tables S1-4**).

### 2.2 Extended Network Diffusion Model (eNDM)

The spread of disease-causing pathological protein species through the brain’s network (connectivity matrix, *C*) over time *t* can be modeled as a diffusion process starting from a seed location, using the Network Diffusion Model (NDM) introduced by Raj *et al*^17^. The NDM assumes that pathology transmission from region 1 to region 2 is a first-order diffusion process along the fiber projections, such that:

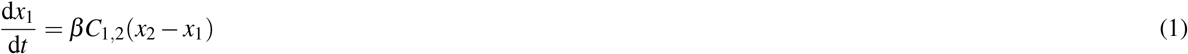

where *x*_1_ and *x*_2_ are the magnitudes of disease-causing pathology in each region and β is a global diffusivity constant, which is considered as the rate of disease progression.

Denoting pathology from all regions *i* into a vector **x**(*t*) = {*x*_*i*_(*t*) ∀ *i* ∈ [1, *N*]}, the above equation becomes:

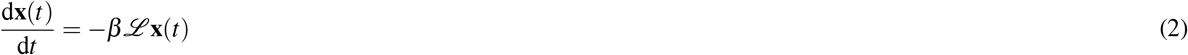

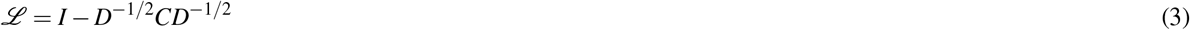

where ℒ is the graph Laplacian and *D* is a diagonal matrix containing the degree (total outgoing connection) of each node. **Equation 2** allows a closed-form solution **x**(*t*) = *e*^−βℒ*t*^**x**_0_ where **x**_0_ is the initial pattern of the disease process at *t* = 0. The matrix exponential *e*^−βℒ*t t*^, the diffusion kernel, acts as a spatial and temporal blurring operator on **x**_0_. The model’s diffusion time *t* is in arbitrary units (a.u.).

In this study, we extended this basic model by adding two key processes: (1) protein aggregation and/or clearance modeled via first-order kinetics, and (2) a continuous seeding process at a prescribed seed site, modeled by a function in time *f*_*τ*_ (*t*). This yielded an extended NDM (“eNDM”), such that:

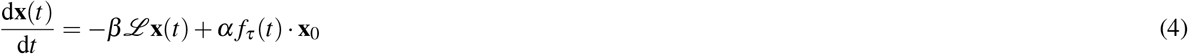

where *α* is a kinetic rate parameter that controls how fast tau is produced by the seeding and templated corruption of healthy tau in the pre-specified seed region **x**_0_. In this study, we chose the bilateral EC as the seeding site due to the proven origin of pathological tau spread in AD from the EC^1^. The function *f*_*τ*_ (*t*) is a continuous-time function that seeks to model the initial rise in seeding activity (“nucleation”), its sustenance over time, followed by its eventual decline. In this work, we impose a simple Gamma shape to this function since the entire function can be parameterized by a single time constant *λ* in the same units as model time *t*, yielding: 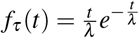.

### 2.3 Model fitting, testing and statistical analysis

#### Model implementation

Unlike the previous uses of the NDM, where a closed-form solution is easy to obtain, there is no closed-form solution to the eNDM (**Equation 4**) due to the additional kinetic terms and a continuous seeding function in time. The theoretical model’s differential equations were therefore numerically solved using MATLAB’s `ode45()` solver.

#### Model inference

The theoretical tau model was fitted to and tested against both individual patients’ re-scaled z-scores and group-averaged regional tau t-statistics, both with respect to matched healthy controls. We chose to only use cross-sectional data at baseline scans for testing because patients would have already accumulated and established pathology at the first imaging. Empirical tau-PET regional vectors were obtained from individual ADNI patients as described above. Regional empirical tau pathology vector **x**_*emp*_ is expressed as 86 × 1 vector corresponding to the 86-region DK brain atlas. Due to the known off-target binding behavior of the tau PET tracer to striatal regions^42^, data from these regions were excluded from our analyses. We also excluded the cerebellar regions, as the cerebellum has long been thought to be relatively preserved in AD^1, 43^ and therefore tau-PET data are often normalized to cerebellar values^43^, as we have done here. Thus, we excluded a total of 10 regions including striatal and cerebellar regions to yield a 76-region tau vector.

To fit the model to the above empirical data, we need to estimate subject-specific optimal model parameters. Recall the eNDM has two rate parameters *α*, β, and a seeding function time scale *λ*, collectively denoted by the (global) parameter set *θ*. We solve the following minimization problem:

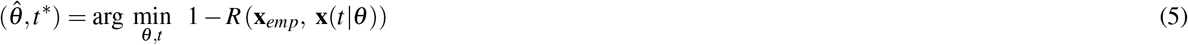

where *R*(·, ·) denotes the Pearson correlation strength between two vector quantities, and *t*^*\**^ represents the model times *t* at which the maximum *R* was achieved. We also experimented with additional prior terms imposing a penalty for high values of *θ* ; however, this option was discarded when we observed almost no significant change in the fits.

#### Model testing and validation

The optimally fitted model was then statistically tested against empirical tau data vectors at both individual and group levels. The primary test statistic used to evaluate all models was Pearson’s correlation coefficient between the empirical and optimally-fitted model: 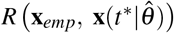, as in **Equation 5**. In each case, the dependent variable was the vector of regional tau levels and the independent variable was the eNDM-predicted regional tau vector. The gene expression vector was used to correlate with observed or residual tau values obtained after regression.

Linear regression was conducted using MATLAB’s `fitlm` function to obtain residual tau values between observed and eNDM-modeled tau (**Figure 1**). In our hypothesis-driven analysis, we aimed to quantify the spatial correspondence between AD-typical imaging signatures of tauopathy and regional expression levels of selected AD-related genes. Univariate correlation between the t-statistics obtained from Student’s t-tests on the regional levels of observed as well as residual tau and regional gene expression levels of the selected genes was quantified using the Pearson correlation coefficient *R*. This allowed us to categorize the genes into those contributing to selective vulnerability (SV) or selective resilience (SR) based on the sign of the correlation between gene expression and observed and residual tau. Principal component analysis (PCA) as well as clustering were conducted on the genes to confirm the gene sets. For qualitative assessment, we illustrate the spread of tau pathology in the three patient groups using brain renderings generated using BrainNet Viewer (http://www.nitrc.org/projects/bnv/)^44^.

### 2.4 Defining four selective vulnerability groupings of genes

The mean Pearson correlation values between regional gene expression and observed and residual tau levels across all patients were used to categorize the genes into the four sets: (1) selective vulnerability genes, network-related (SV-NR), (2) selective vulnerability genes, network-independent (SV-NI), (3) selective resilience genes, network-related (SR-NR), (4) selective resilience genes, network-independent (SR-NI). Briefly, genes implicated in vulnerability were those positively correlated with observed and residual tau, whereas genes implicated in resilience were those with negative correlation. Whenever the correlation with observed tau was higher than residual tau, the gene function was categorized to be related to the brain’s connectome (“network-related”, NR). On the other hand, a higher correlation of the gene with residual than observed tau implicated its function to be independent of the brain’s connectome (“network-independent”, NI). These four sets with example genes from each of them are illustrated using the violin plots in Figure 3.

### 2.5 Gene ontology analyses

We conducted separate gene ontology (GO) analyses on the four gene sets by employing the ShinyGO toolbox (version 0.77, http://bioinformatics.sdstate.edu/go/)^45^ to identify functional enrichment patterns and their associated gene networks. We chose the GO biological processes database, used an FDR cutoff of 0.05, sorted the functions based on fold enrichment of the genes, and used the gene counts from the GO biological processes database for background. Dot-plot charts and networks were constructed to visualize the functional networks of the four gene sets.

## 3 Results

### 3.1 eNDM accurately predicts the progression of tau pathology across patient cohorts

**Figure 2A&B** illustrate the regional distribution of observed (AV1451-PET-derived) and extended Network Diffusion Model (eNDM)-predicted tau pathology, respectively. The regional data were obtained using the pre-processing volumetric pipeline as described in **Methods** on the 86-region DK atlas parcellation for 196 subjects diagnosed with EMCI, LMCI, or AD. The eNDM was evaluated at the group-level 86 × 86 structural connectome acquired from healthy brains from the HCP^28^. To minimize non-pertinent sources of variability and to ensure statistical interpretability, we fitted the model on all patients using the same canonical pathology seeding site in the EC, but with subject-specific model parameters (see **Supplementary Figure S1** for distribution of the optimal pathology diffusion parameter, β).

**Figure 2:**
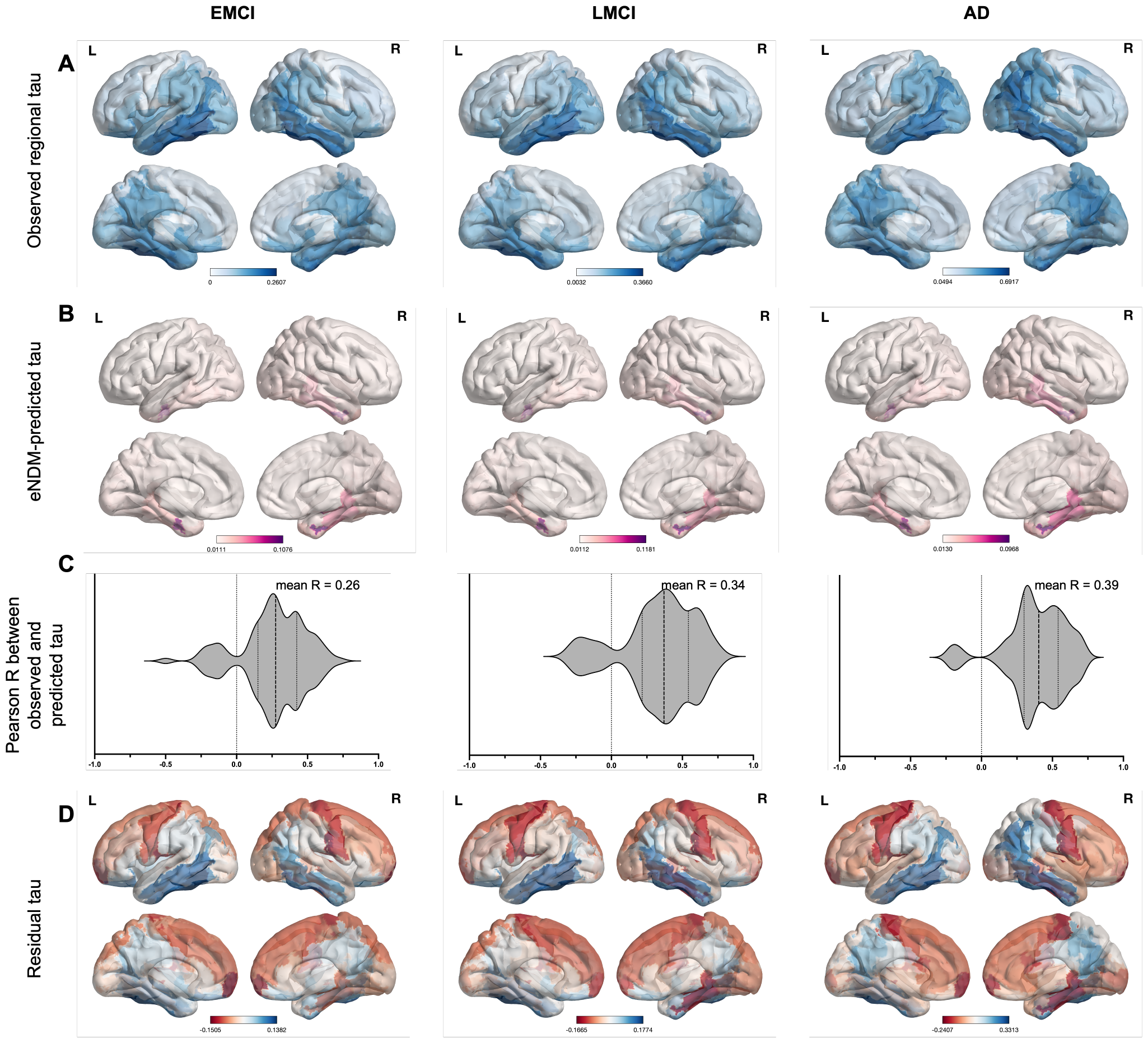
Regional distribution of tau in patients with EMCI, LMCI, and AD. Panel A: Regional distribution of observed (PET-measured) tau levels in the three patient cohorts. Panel B: Regional distribution of extended network diffusion modeled (eNDM-predicted) tau levels in the three cohorts. Panel C: Distribution of the Pearson R correlation values between the observed and predicted tau levels. Panel D: Residual (observed *–* predicted) tau levels in the three groups. Blue and red regions correspond to positive and negative residual tau, respectively.

The spatial correspondence between observed and model-predicted tau pathology is visually apparent in all groups: EMCI, LMCI, and AD. The distribution of Pearson’s correlation (*R*) between each subject’s observed and predicted tau is displayed in **Figure 2C**. The eNDM performed well in all groups, exhibiting statistically significant mean *R* (all *p* < 0.05 by one-sample t-test following Fisher’s *R*-to-z transformation). As expected, eNDM fitted the AD group the best (mean *R* = 0.39 *±* 0.21), followed by LMCI (mean *R* = 0.34 *±* 0.26) and EMCI (mean *R* = 0.26 *±* 0.23). It also successfully recapitulated the classic temporal-limbic-posterior presentation of AD-associated tauopathy. Interestingly, a small subset of patients in each group gave non-significant or even negative *R*, signifying that the model did not work on a small minority of patients. This seems to be a general feature of all theoretical network models, and the observation that this applies to mostly the EMCI and LMCI groups may be expected because not all of these patients may be on the AD spectrum.

Next, we calculated each subject’s and each region’s residual (observed *–* predicted) tau; their group averages are rendered in **Figure 2D**. These residuals were obtained from linear regression fits performed on individual patients as well as group-level averages (see **Methods**). Positive residual indicates that factors above and beyond the connectome-mediated spread contributed to tau progression. We observed positive residual tau in the temporal, posterior parietal, and orbitofrontal regions, in that order, throughout the three groups. Conversely, negative residuals (where the eNDM over-predicted tau pathology) were observed in the fronto-parietal and thalamic regions across all groups, as well as some in the occipital regions of the EMCI and LMCI groups (**Figure 2D**).

### 3.2 AD risk genes show spatial association to tau either directly or after removing network vulnerability

To assess the molecular factors mediating regional SV and SR, we used regional expression data from the AHBA^37^ for 100 selected AD risk genes (**Supplementary Tables 1-4**)^32–36^. Since these data were obtained from healthy donors, these gene regional patterns constitute the *baseline* gene expression and therefore do not consider disease-related changes in transcription. To differentiate between *network-related* (NR) and *network-independent* (NI) mediation of pathology, we calculated the *R* between each gene’s expression pattern and each patient’s observed (*R*_observed_) and residual (*R*_residual_) tau pathology.

From these Tables, we observe that of the 100 genes, 25 gave a strong direct association with regional tau (defined as |*mean R*_*observed*_| ≥ 0.20), confirming that the role of risk genes from GWAS is largely mirrored in their spatial association at baseline with tau. Remarkably, even more risk genes were correlated with residual tau (|*mean R*_*residual*_| ≥ 0.20): 39 out of 100 genes. That far more genes are associated with residual tau than with directly observed tau underscores the necessity of accounting for network effects, and its value in assessing SV/SR of risk genes, many of which would remain undetected through conventional analysis.

### 3.3 AD genes form 4 classes based on their contributions to network-related and network-independent SV and SR

Next we separated the AD risk genes into four classes with respect to two factors: 1) whether the gene confers SV (mean *R*_observed_ > 0) or SR (mean *R*_observed_ < 0); and 2) whether that SV/SR is NR (mean *R*_observed_ > mean *R*_residual_) or NI (mean *R*_residual_ > mean *R*_observed_). Out of the 100 genes, 26 fell into the SV-NR group, 18 in the SV-NI group, 35 in the SR-NR group, and 21 in the SR-NI group. Mean correlations were assessed after combining the AD, LMCI, and EMCI cohorts into one group (*n* = 196). **Figure 3** depicts the distributions of *R* values with respect to observed and residual tau across all patients, separated by the 4 gene classes, for several genes of interest.

**Figure 3:**
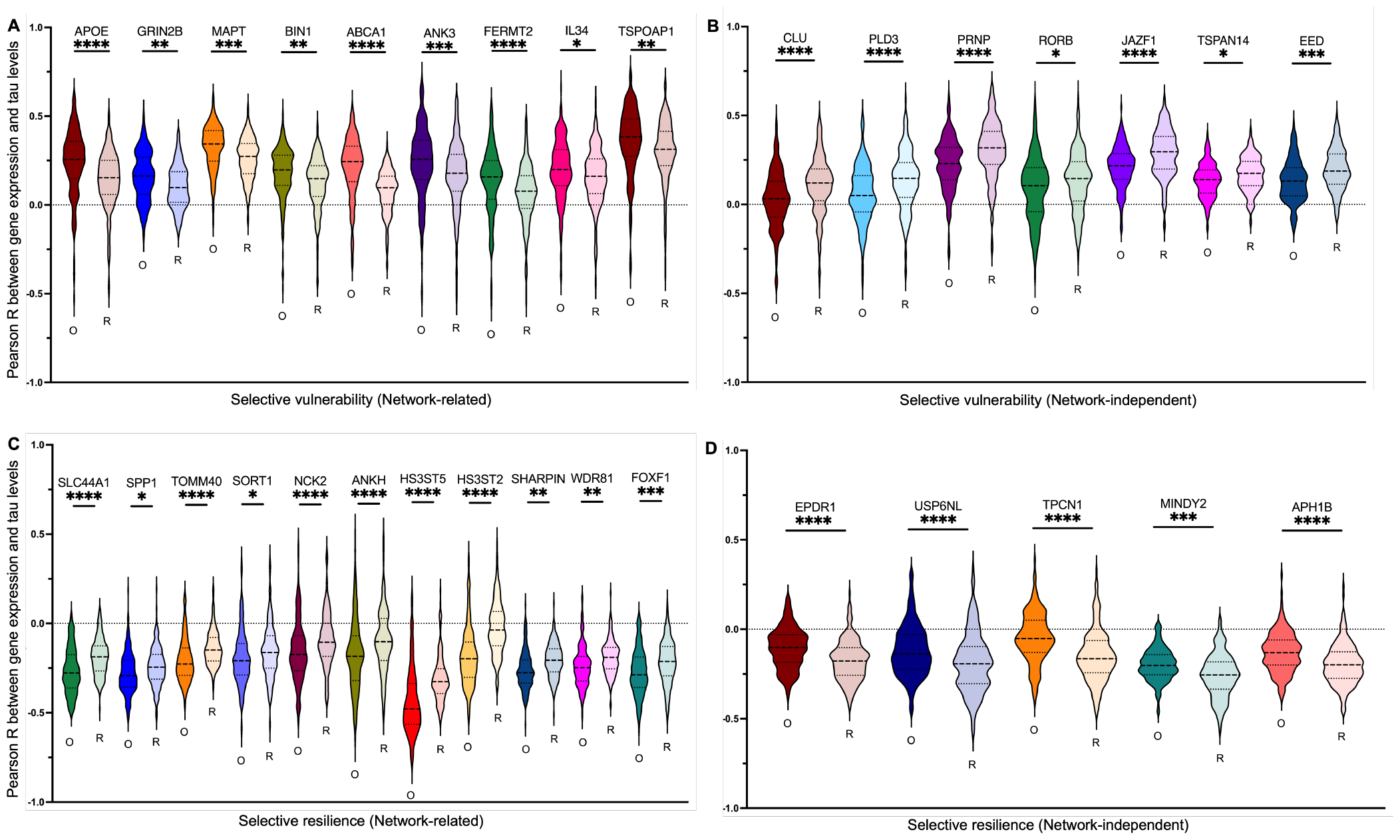
Distribution of Pearson R between gene expression and tau levels across 196 individual patients. Panels A and B include some genes contributing to selective vulnerability towards AD tau pathology, and panels C and D include some genes contributing to selective resilience towards AD tau pathology. Each set further has genes that are either more strongly correlated with observed or residual levels of tau. Each gene has two violin plots associated with it: one with correlations between gene expression and observed tau (marked with an O) and the other with correlations between gene expression and residual tau (marked with an R). Note that the asterisks denote the significance of the comparison between the O and the R violin for each gene: * – *p* < 0.05; ** – *p* < 0.01; *** – *p* < 0.001; **** – *p* < 0.0001.

We found that some of the most well-known SV genes, including *APOE, MAPT* and *BIN1*, were a part of the SV-NR class (**Figure 3A**), while others, such as *PRNP* and *RORB*, belonged to SV-NI class (**Figure 3B**). The latter class is intriguing: its genes are well-known risk factors, yet their direct association with regional tau is low, while their network-decoupled association is high. For the SR genes, *ANKH* and *SORT1*, implicated as protective factors in centenarians, and *NCK2*, a microglial gene downregulated in *APOE* ε4/ε4 patients, belonged to the SR-NR class. *EPDR*, another protective factor in centenarians, belonged to the SR-NI class. We further discuss the implications of NR and NI vulnerability and resilience in the **Discussion**.

### 3.4 Gene classes at the group level

To investigate the gene classes identified through the individual-level analysis above, we also examined their relationships at a group level. On t-statistics per region across the 196 patients for observed and residual tau, we calculated its Pearson *R* against each gene’s regional expression. **Figure 4A** shows *R*_observed_ plotted against *R*_residual_, where each point represents a single gene color-coded by its class identity: red (SV-NR), magenta (SV-NI), blue (SR-NR), and cyan (SR-NI). We observed distinct segregation based on whether gene expression correlated positively or negatively with observed or residual tau (**Figure 4A**). We found that there was a strong association between these two measures (*R*^2^ = 0.53, *p* < 10^−6^), implying that genes tended to contribute to both direct as well as indirect vulnerability to tau after accounting for the underlying network transmission process. We further note that the best-fit line found by linear regression had a slope of 0.60 (solid line) [95 % CI = 0.48, 0.71], which was significantly less than 1 (dashed line; *p* < 0.001). This indicates that genes generally exhibited higher correlations to residual rather than observed tau, suggesting that they contribute less to the direct vulnerability of regions than to the additional vulnerability that cannot be explained by the canonical, network-based spread of tau.

**Figure 4:**
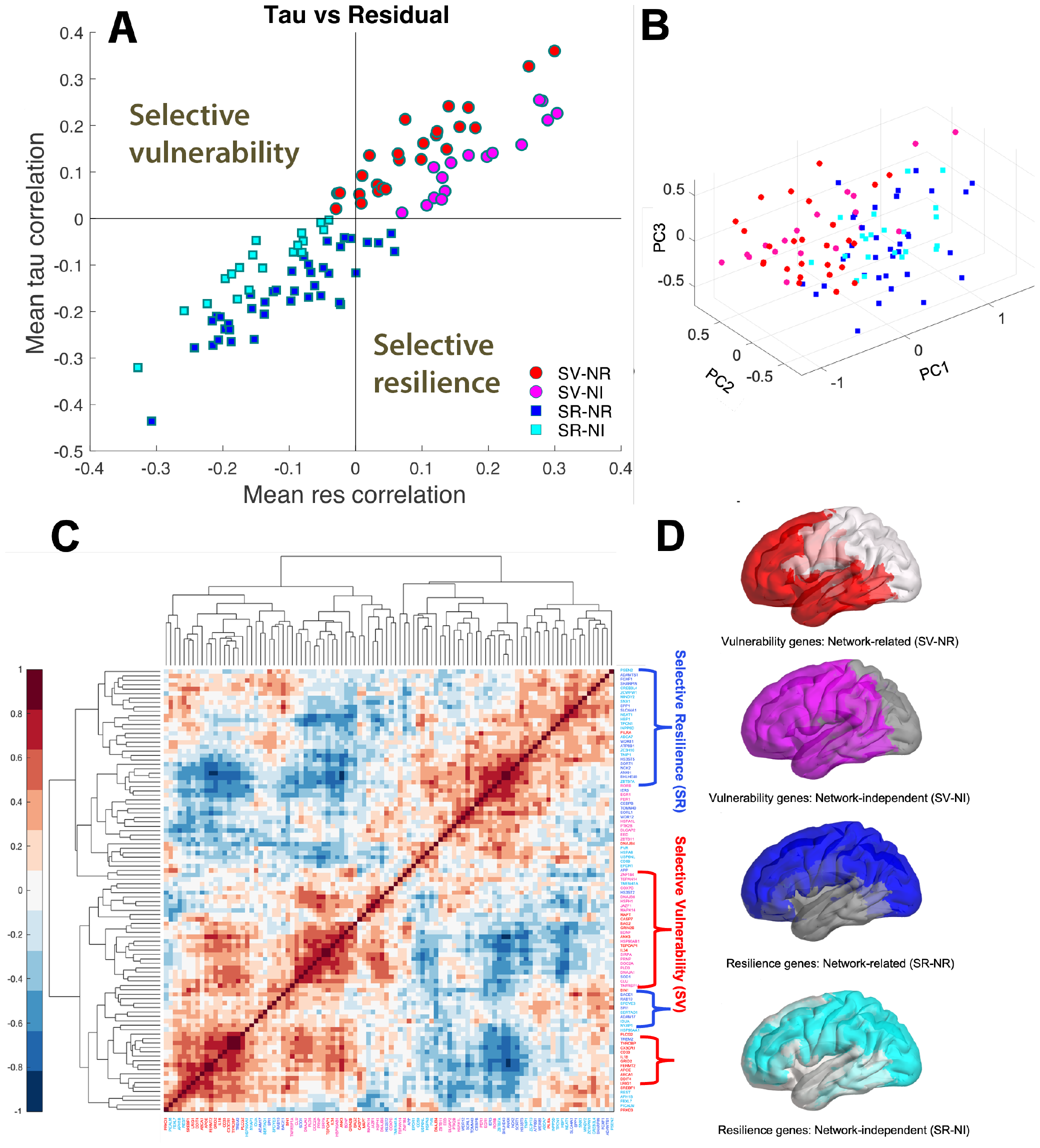
Group level analysis of genes implićated in Alzheimer’s disease using t-scores. AD ene classes separated with respect to two factors: 1) whether the gene confers SV (mean *R*_observed_ > 0) or SR (mean *R*_observed_ < 0); and 2) whether that SV/SR is NR (mean *R*_observed_ > mean *R*_residual_) or NI (mean *R*_residual_ > mean *R*_observed_). Out of the 100 genes, 26 fell into the SV-NR group, 18 in the SV-NI group, 35 in the SR-NR group, and 21 in the SR-NI group. Mean correlations were assessed after combining the AD, LMCI, and EMCI cohorts into one group (*n* = 196). See tables 5-8 in supplemental

#### Spatial clustering of gene classes

We next asked whether this four-way classification was apparent in the clustering of their spatial profiles. PCA was performed on the genes’ spatial distributions: the first 3 PCs are shown in **Figure 4B** for all 100 genes, which we color-coded following the same conventions as in **Figure 4A**. Overall, the 4 classes formed distinct clusters in PC space, indicating that genes within a class exhibited more similarity to each other than to those in the other classes. We attempted to confirm these findings using hierarchical clustering (**Figure 4C**), which also indicated that the SV and SR gene sets generally tended to group together. Clustering was less distinct between NR and NI genes, but we also found several smaller clusters containing single subgroups (**Figure 4C**). Thus, we conclude that genes that have similar spatial patterns also tend to have a similar role in mediating SV/SR to human AD-related tauopathy.

### 3.5 SV-NR, SV-NI, SR-NR, and SR-NI genes exhibit distinct spatial patterns

To understand the spatial distributions of the 4 gene classes at the aggregate level, we performed PCA on their regional expression and rendered each class’s first principal component (PC1) in **Figure 5** (for illustration purposes these spatial patterns were thresholded such that only the highest half of regional density is shown). The visual findings are both expected (medial and temporal dominance of vulnerable genes vs occipital dominance of resilient genes) and unexpected (high density in frontal areas of vulnerable genes and parietal areas of resilient genes). The PC1 patterns continue to show the double-dissociation suggested earlier: correlation between PC1 of SV-NR genes was significantly positive with tau and non-significant with residual tau, whereas that of SV-NI genes had the reverse effect. Analogously for the SR genes. We therefore conclude that these four sets of genes constitute spatially distinct classes.

**Figure 5:**
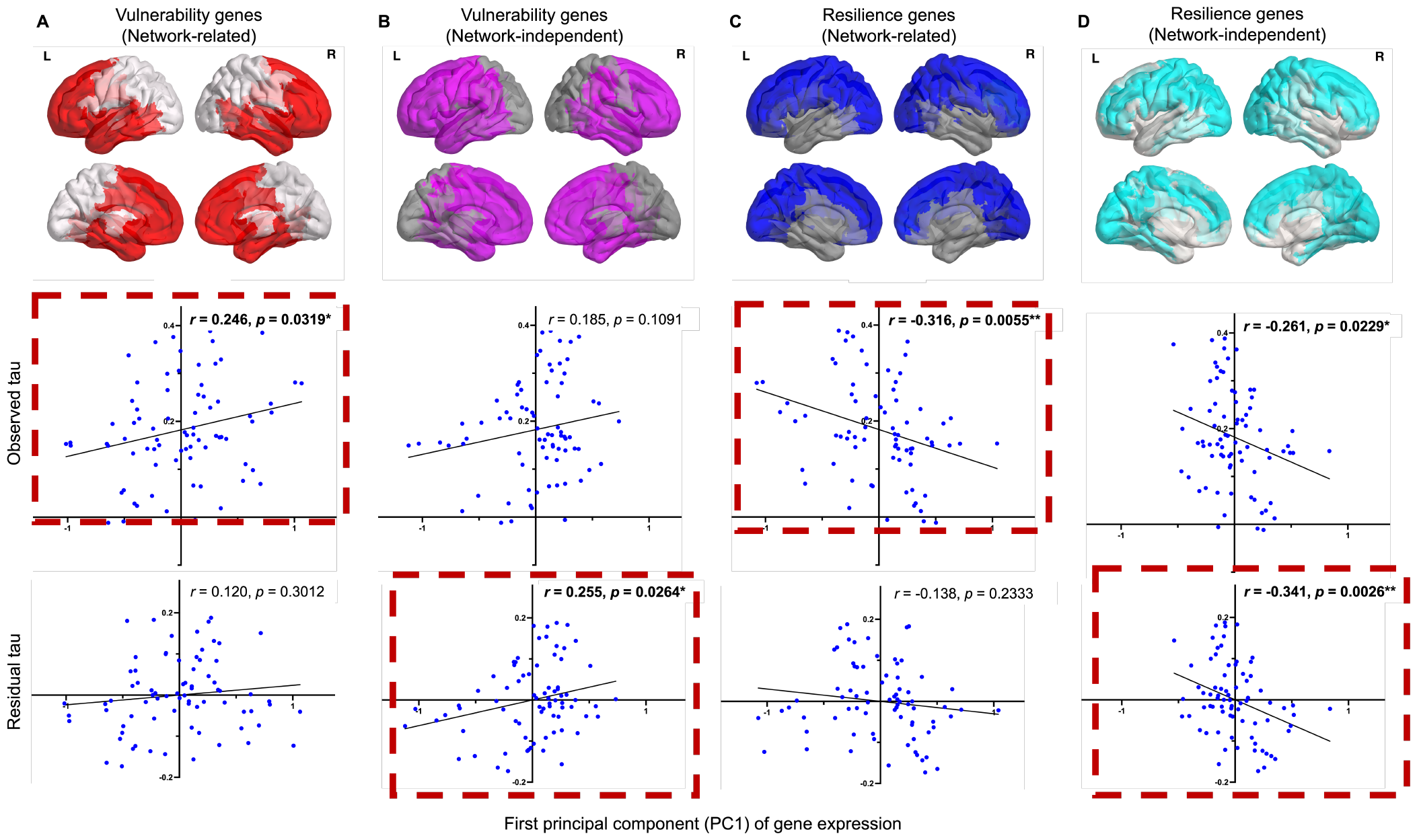
Correlation between the first principal component (PC1) of gene expression and tau levels across brain regions. Distributions of PC1 of the gene sets contributing to selective vulnerability (panels A and B) and selective resilience (panels C and D) towards AD pathology in 76 brain regions are depicted as brain renderings. PC1 of the vulnerability group is positively correlated with observed as well as residual tau (dashed red squares around plots in panels A and B), whereas that of the resilience groups shows a negative correlation to observed and residual tau (dashed red squares around plots in panels C and D). Significant correlations are indicated by asterisks as well as bolded *R* and *p* values. The color coding is maintained as red – vulnerability (network-related); magenta – vulnerability (network-independent); blue – resilience (network-related); cyan – resilience (network-independent). Signficance cutoffs: * – *p* < 0.05; ** – *p* < 0.01.

### 3.6 Gene ontology analysis reveals distinct functional networks for each gene class

We hypothesized that, if genes can impart differential vulnerability and resilience in NR or NI manners, then distinct biological processes may be associated with them. We used the ShinyGO tool^45^ to examine the functional enrichment for each gene class using gene ontology (GO) analysis. **Figure 6** shows dot plots of the biological processes with the highest fold enrichment for the SV-NR, SV-NI, SR-NR, and SR-NI classes, respectively. The network relationship between these processes is also shown.

**Figure 6:**
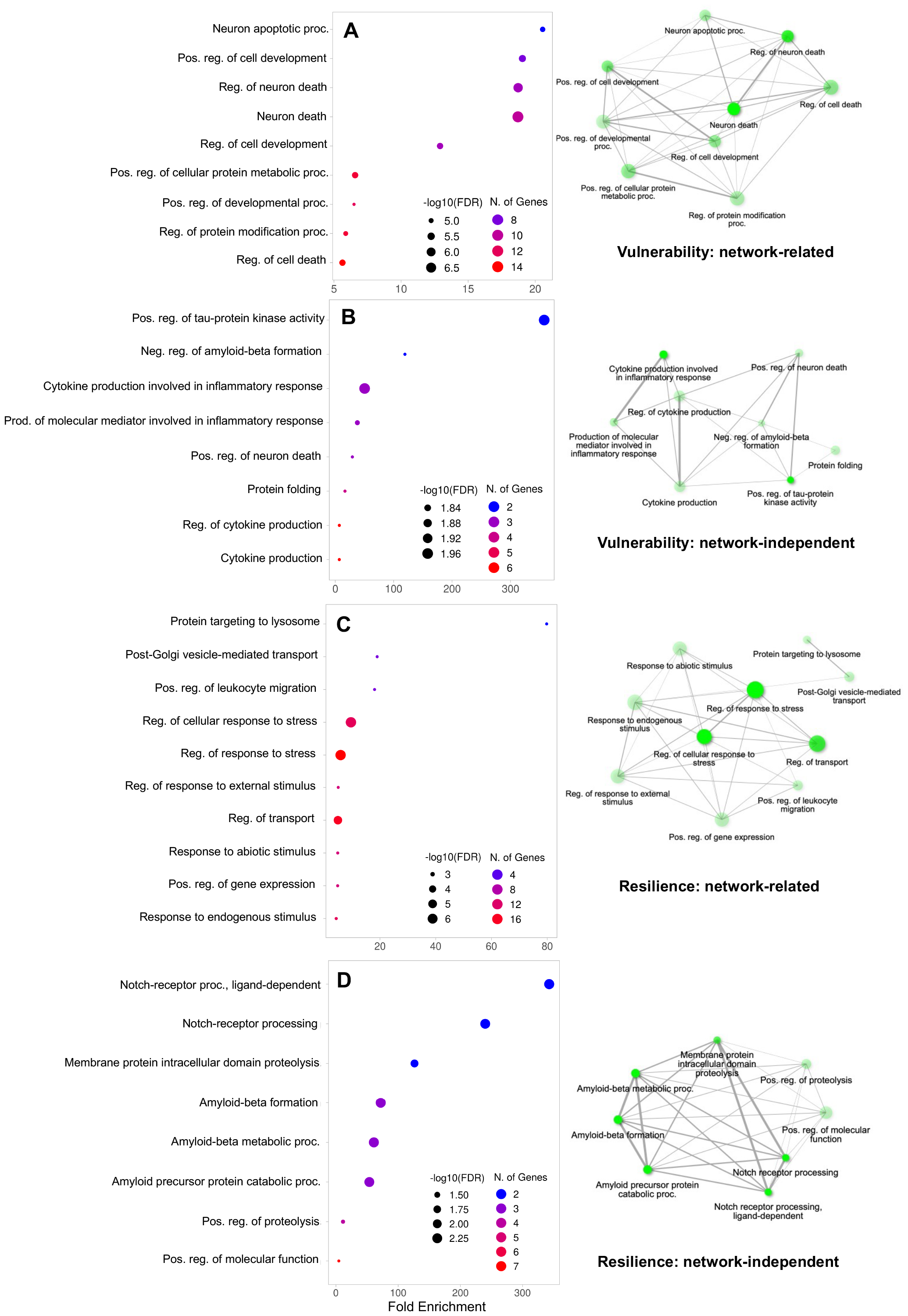
Functional enrichment networks of the four gene sets. Panels display a maximum of 10 GO biological processes surviving an FDR cutoff of 0.05 and sorted based on fold enrichment for genes belonging to the selective vulnerability (A and B) and the selective resilience (C and D) sets, respectively. These results indicate that the 4 broad vulnerability classes have divergent functional roles and are embedded in different functional networks with very different biological roles.

We found that the functions of each set of genes were remarkably dissimilar to each other. The SV-NR subgroup was enriched in genes relating to developmental processes and in particular apoptosis (**Figure 6A**). Notably, all of these processes are highly linked to each other, suggesting that these genes can be grouped in the same functional network. The SV-NI genes were predominantly involved in inflammation as well as amyloidosis, tau misfolding, and neuron death, which are distinct but related processes (**Figure 6B**). Conversely, we found that the functions of the SR-NR genes were all related to cellular responses to stress (**Figure 6C**), while SR-NI genes predominantly contributed to proteolysis and Aβ formation in particular (**Figure 6D**). The specific biological processes implicated using GO analysis were themselves tightly functionally related, as seen in the accompanying networks.

## 4 Discussion

The question of why certain regions, structures, cells, or systems become selectively vulnerable (SV) or remain resilient (SR) to neurodegenerative pathology is one of the most important open questions in the field. While ramifying AD pathology can and will cause alterations in the expression of risk genes, a related but separate question that has received less attention is whether and how the presence of certain molecular factors and their gene regulators at *baseline* (that is, in the non-disease condition) may confer inherent vulnerability or resilience even before the establishment of overt pathology. A full exposition of candidate genes and networks behind SV/SR may help the field of AD in its quest to understand mechanisms, find effective drugs, and translate druggable targets to patients.

A key challenge in understanding the molecular underpinnings of this innate SV or SR is that vulnerability and the ensuing pathology are likely not just the result of a few genes, pathways, or neural systems acting in isolation. Indeed, there is a well-known and puzzling spatial dissociation between upstream risk genes and downstream disease topography^13, 14^. Hence these questions must be assessed quantitatively via computational models, on all risk genes, on the whole brain and across many individuals. In this study, we used a combination of mathematical modeling and comprehensive transcriptomic data to explore the underpinnings of SV and SR in the context of AD pathology. We sought to understand how *innate* SV/SR is associated with regional gene expression, in relation to the effect of the network-based spread of tau.

### 4.1 Summary of major findings

We first demonstrated that the extended Network Diffusion Model, based on (eNDM)^17, 21^, highly significantly predicted regional tau-PET signal in the vast majority of EMCI, LMCI and AD patients in the ADNI database (**Figure 2C**). However, the eNDM prediction was not complete, systematically underestimating tau in limbic and temporal areas and over-predicting it elsewhere (**Figure 2D**). Armed with this network vulnerability model, we reassessed the effect of top 100 AD risk factor genes, looking specifically for whether their association with tau is direct or indirect through the residual tau (**Figure 3**).

We hypothesized that AD risk genes could contribute to innate SV or SR in one of two ways: 1) ‘network-related’ (NR), where the gene was more correlated to observed tau pathology than to the “residual” tau pathology after removing network-based spread effects; or 2) ‘network-independent’ (NI), where the gene was more correlated to the residual than the observed tau. In this manner, we were able to use the eNDM to separate out genes based on whether their effects were more coupled to the network transmission process (SV-NR or SR-NR) or to processes unrelated to network spread (SV-NI or SR-NI).

We found that out of the 100 selected AD risk genes, 26 belonged to the SV-NR class, 18 to the SV-NI class, 35 to the SR-NR class, and 21 to the SR-NI class. Certain genes, such as *PRNP, JAZF1* and *RORB*, are well-known risk factors, yet their direct association with regional tau is low, while their network-decoupled association is high. Indeed, risk genes overall showed significantly higher association to residual than to observed tau, group level (**Figure 4A**) and at the individual level where 25 genes were strongly correlated with tau directly, while 39 genes were correlated to residual tau (**Supplementary Tables 1-4**). This could be a potential explanation for the puzzling dissociation between upstream genes and downstream pathology^**?**,**?**^.

The genes in each class tended to cluster together based on their expression patterns (**Figure 4B&C**). Overall, each class exhibited distinct, canonical spatial distributions (**Figure 4D**) that show double dissociation with observed vs residual tau (**Figure 5**). These 4 SV/SR classes, encompassing both network-related and -independent modes of action, can help us understand how innate vulnerability plays through genes and how it interacts with network-mediated transmission of pathology.

Finally, we asked whether the dissociations found between the 4 classes are underpinned by their divergent functional roles. Our GO analysis (**Figure 6**) identified distinct functional enrichments and networks for each of these gene sets. The SV-NR genes had functions closely linked to neuronal death, whereas the SV-NI ones pointed to functions involved in cytokine regulation, kinase activity, and the overall inflammatory response. On the other hand, the SR-NR group was implicated in immune response and transport functions, with the SR-NI group functioning more directly with proteolysis and Aβ metabolism. Overall, these results suggest that SV-NR, SV-NI, SR-NR, and SR-NI are mediated by genes that are distinct not only spatially but also functionally. The genes in the four broad classes constitute diverse genetic, molecular, and biological functions, and have equally diverse distributions in the brain.

Let us now briefly assess the known physiological and pathological processes regulated by the most prominent genes from each vulnerability class.

### 4.2 Genes implicated in selective vulnerability to AD

The strongest network-related association with tau vulnerability was observed in *APOE, MAPT* and *TSPOAP1. APOE* is one of the most well-known genetic risk factors for AD, whose ε4 allele is well known to confer a higher risk and reduce the age of onset of the disease. *APOE* is normally involved in lipid and cholesterol metabolism, but in AD, *APOE* binds to Aβand affects the clearance of soluble Aβ. The ε4 allele specifically increases the rate of Aβ aggregation^32^ and the spreading of tau downstream of amyloidosis^34^. The *MAPT* gene encodes tau protein which stabilizes microtubules, but in AD, tau hyperphosphorylates, aggregates, and spreads throughout the brain trans-neuronally^18, 46^, resulting in neurodegeneration. Polymorphisms associated with *MAPT* increase AD risk and predispose tau to misfold and aggregate. *TSPOAP1* encodes a cytoplasmic protein closely associated with its mitochondrial transmembrane protein partner translocator protein (TSPO), and is involved in neuroinflammatory processes^**?**^. It shows one of the strongest direct signatures of selective vulnerability in our study.

The strongest network-independent vulnerability to tau was displayed by *CLU, PRNP* and *JAZF1. CLU*, the clusterin gene, is the third most significant genetic risk factor for AD, whose variants have been linked to altered cognitive and memory function and brain structure. Clusterin is a multifunctional glycoprotein involved in lipid transport and immune modulation, as well as in pathways of cell death and survival, oxidative and proteotoxic stress^47^. The *PRNP* gene encodes prion protein (PrP), implicated in Creutzfeldt-Jakob disease, a canonical example of trans-neuronally transmitted disease^48^. *JAZF1* is implicated in type-2 diabetes (T2D) in addition to AD, and our analysis showed that its regional expression levels were highly correlated with residual levels of tau, suggesting its plausible function as an AD risk factor. Due to the link between T2D and AD^49^, *JAZF1* poses an excellent candidate gene open to further exploration as an AD risk gene.

### 4.3 Genes implicated in selective resilience to AD

The genes that we identified as contributing to SR are enriched in endo-lysosomal and immune functions as well as amyloid beta formation and clearance (**Figure 6C&D**). Among the SR genes, *SLC44A1* codes for a plasma membrane choline transporter, essential for the synthesis of acetylcholine (ACh), is downregulated in the microglia of AD brains^50^. The *TOMM40* gene encodes the outer mitochondrial membrane pore subunit necessary for oxidative phosphorylation and enables protein transport into mitochondria^51^. GWAS have reported an increased risk for AD associated with the rs2075650 SNP of *TOMM40* minor allele^52^. *USP6NL* is a GTPase-activating protein involved in the control of endocytosis. It is implicated in the dysfunction of the myeloid endolysosomal system in AD^53^. *ANKH, SORT1, EPDR1* were identified in a recent GWAS of AD and aging^54^. *SORT1* is involved in endosomal trafficking. *EPDR1* plays a crucial role in the adhesion of neural cells and dopaminergic regulation of neurogenesis and is downregulated in AD. *TPCN1*, another SR-related gene, codes for endosomal cation-permeable channels.

### 4.4 Limitations

An important limitation of our study is that while high-quality tau-PET patient data are available from ADNI, gene expression data from AHBA are from different and far fewer healthy subjects. This was appropriate for the current purpose of exploring baseline genetic markers of AD, but future studies on disease modification of gene expression would require spatial transcriptomic data in AD individuals – an effort that is ongoing in the field. Further, gene samples in AHBA are sparse, sufficient for current regional analysis, but not extensible to finer resolutions.

### 4.5 Conclusions

We conclude from these findings that risk genes in AD are associated at baseline with the spatial patterning of tau in subsequent disease. However, their association is not entirely or even mainly with observed tau, but in a joint interaction with network vulnerability. After accounting for the network interaction, we were able to characterize the role of genes in governing innate regional SV and SR. We identified 4 gene classes, which display distinct and segregated functional roles. Together these data throw new light on the genetic and network underpinnings of selective vulnerability and resilience in AD. Identification of disease-modifying gene networks in the healthy brain may be crucial to designing targeted therapeutic manipulation in early disease stages.

## 5 Data availability

We plan to share all relevant data (regional gene expression, tau) and source code publicly upon paper acceptance, via our GitHub repository (https://github.com/Raj-Lab-UCSF). Original MRI and PET images may be obtained directly from ADNI.

## Acknowledgements

We thank ADNI and the AHBA for making their data available to us. Thanks also to Benjamin Sipes at UCSF for his assistance.

## Funding

This study was partially supported by NIH grants R01NS092802, RF1AG062196, and R01AG072753 awarded to Ashish Raj.

## Competing interests

The authors have no competing financial or other interests.

## Author contributions statement

A.R. conceived the study. C.A. conducted the literature search, analyzed the data, and wrote the manuscript. J.T. helped with data analysis. J.T. and P.D.M. helped with the generation of brain figures and the implementation of underlying algorithms. All authors reviewed the manuscript.

## Notes

### Competing Interest Statement

The authors have declared no competing interest.

## References

1. Braak, H. & Braak, E. Neuropathological stageing of alzheimer-related changes. Acta Neuropathol 82, 239–59 (1991).

2. Seeley, W. W. Selective functional, regional, and neuronal vulnerability in frontotemporal dementia. Curr Opin Neurol 21, 701–7, DOI: 10.1097/WCO.0b013e3283168e2d (2008).

3. Fu, H., Hardy, J. & Duff, K. E. Selective vulnerability in neurodegenerative diseases. Nat Neurosci 21, 1350–1358, DOI: 10.1038/s41593-018-0221-2 (2018).

4. Saxena, S. & Caroni, P. Selective neuronal vulnerability in neurodegenerative diseases: from stressor thresholds to degeneration. Neuron 71, 35–48, DOI: 10.1016/j.neuron.2011.06.031 (2011).

5. Roussarie, J. P. et al. Selective neuronal vulnerability in alzheimer’s disease: A network-based analysis. Neuron 107, 821–835 e12, DOI: 10.1016/j.neuron.2020.06.010 (2020).

6. Saxena, S. & Caroni, P. Selective Neuronal Vulnerability in Neurodegenerative Diseases: from Stressor Thresholds to Degeneration. Neuron 71, 35–48, DOI: 10.1016/j.neuron.2011.06.031 (2011).

7. Wang, M. et al. Integrative network analysis of nineteen brain regions identifies molecular signatures and networks underlying selective regional vulnerability to Alzheimer’s disease. Genome Medicine 8, 104, DOI: 10.1186/s13073-016-0355-3 (2016).

8. Grothe, M. J. et al. Molecular properties underlying regional vulnerability to alzheimer’s disease pathology. Brain 141, 2755–2771, DOI: 10.1093/brain/awy189 (2018).

9. Kunkle, B. W. et al. Genetic meta-analysis of diagnosed alzheimer’s disease identifies new risk loci and implicates abeta, tau, immunity and lipid processing. Nat Genet. 51, 414–430, DOI: 10.1038/s41588-019-0358-2 (2019).

10. Mezias, C., LoCastro, E., Xia, C. & Raj, A. Connectivity, not region-intrinsic properties, predicts regional vulnerability to progressive tau pathology in mouse models of disease. Acta Neuropathol Commun 5, 61 (2017).

11. Torok, J., Maia, P. D., Verma, P., Mezias, C. & Raj, A. Emergence of directional bias in tau deposition from axonal transport dynamics. PLoS Comput. Biol 17, e1009258, DOI: 10.1371/journal.pcbi.1009258 (2021).

12. Raj, A. Graph models of pathology spread in alzheimer’s disease: An alternative to conventional graph theoretic analysis. Brain Connect. 11, 799–814, DOI: 10.1089/brain.2020.0905 (2021).

13. Subramaniam, S. Selective neuronal death in neurodegenerative diseases: The ongoing mystery. Yale J Biol Med 92, 695–705 (2019).

14. Fusco, F. R. et al. Cellular localization of huntingtin in striatal and cortical neurons in rats: lack of correlation with neuronal vulnerability in huntington’s disease. J Neurosci 19, 1189–202, DOI: 10.1523/JNEUROSCI.19-04-01189.1999 (1999).

15. Chung, C. G., Lee, H. & Lee, S. B. Mechanisms of protein toxicity in neurodegenerative diseases. Cell Mol Life Sci 75, 3159–3180, DOI: 10.1007/s00018-018-2854-4 (2018).

16. Kaufman, S. K. et al. Tau prion strains dictate patterns of cell pathology, progression rate, and regional vulnerability in vivo. Neuron 92, 796–812, DOI: 10.1016/j.neuron.2016.09.055 (2016).

17. Raj, A., Kuceyeski, A. & Weiner, M. A network diffusion model of disease progression in dementia. Neuron 73, 1204–15 (2012).

18. Clavaguera, F. et al. Transmission and spreading of tauopathy in transgenic mouse brain. Nat. Cell Biol. 11, 909–913, DOI: 10.1038/ncb1901 (2009).

19. Iba, M. et al. Synthetic tau fibrils mediate transmission of neurofibrillary tangles in a transgenic mouse model of alzheimer’s-like tauopathy. J Neurosci 33, 1024–37 (2013).

20. Raj, A. et al. Network diffusion model of progression predicts longitudinal patterns of atrophy and metabolism in alzheimer’s disease. Cell Rep 10, 359–369 (2015).

21. Anand, C., Maia, P. D., Torok, J., Mezias, C. & Raj, A. The effects of microglia on tauopathy progression can be quantified using nexopathy in silico (nexis) models. Sci Rep 12, 21170, DOI: 10.1038/s41598-022-25131-3 (2022).

22. Warren, J. D. et al. Molecular nexopathies: a new paradigm of neurodegenerative disease. Trends Neurosci 36, 561–9, DOI: 10.1016/j.tins.2013.06.007 (2013).

23. Seeley, W. W., Crawford, R. K., Zhou, J., Miller, B. L. & Greicius, M. D. Neurodegenerative diseases target large-scale human brain networks. Neuron 62, 42–52, DOI: 10.1016/j.neuron.2009.03.024 (2009).

24. Desikan, R. S. et al. An automated labeling system for subdividing the human cerebral cortex on mri scans into gyral based regions of interest. Neuroimage 31, 968–80, DOI: 10.1016/j.neuroimage.2006.01.021 (2006).

25. Gorgolewski, K. et al. Nipype: a flexible, lightweight and extensible neuroimaging data processing framework in python. Front Neuroinform 5, 13, DOI: 10.3389/fninf.2011.00013 (2011).

26. Jenkinson, M., Bannister, P., Brady, M. & Smith, S. Improved optimization for the robust and accurate linear registration and motion correction of brain images. Neuroimage 17, 825–41, DOI: 10.1016/s1053-8119(02)91132-8 (2002).

27. Dayan, M. et al. Profilometry: A new statistical framework for the characterization of white matter pathways, with application to multiple sclerosis. Hum Brain Mapp 37, 989–1004, DOI: 10.1002/hbm.23082 (2016).

28. McNab, J. A. et al. The human connectome project and beyond: initial applications of 300 mt/m gradients. Neuroimage 80, 234–45, DOI: 10.1016/j.neuroimage.2013.05.074 (2013).

29. Abdelnour, F., Voss, H. U. & Raj, A. Network diffusion accurately models the relationship between structural and functional brain connectivity networks. Neuroimage 90, 335–47, DOI: 10.1016/j.neuroimage.2013.12.039 (2014).

30. Owen, J. P. et al. The structural connectome of the human brain in agenesis of the corpus callosum. Neuroimage 70, 340–55, DOI: 10.1016/j.neuroimage.2012.12.031 (2013).

31. Verma, P., Nagarajan, S. & Raj, A. Spectral graph theory of brain oscillations–revisited and improved. Neuroimage 249, 118919, DOI: 10.1016/j.neuroimage.2022.118919 (2022).

32. Karch, C. M., Cruchaga, C. & Goate, A. M. Alzheimer’s disease genetics: from the bench to the clinic. Neuron 83, 11–26, DOI: 10.1016/j.neuron.2014.05.041 (2014).

33. Ayoub, C. A., Wagner, C. S. & Kuret, J. Identification of gene networks mediating regional resistance to tauopathy in late-onset alzheimer’s disease. PLoS Genet. 19, e1010681, DOI: 10.1371/journal.pgen.1010681 (2023).

34. Seto, M., Weiner, R. L., Dumitrescu, L. & Hohman, T. Protective genes and pathways in alzheimer’s disease: moving towards precision interventions. Mol Neurodegener 16, 16, DOI: 10.1186/s13024-021-00452-5 (2021).

35. Andrews, S. J., Fulton-Howard, B. & Goate, A. Interpretation of risk loci from genome-wide association studies of alzheimer’s disease. Lancet Neurol 19, 326–335, DOI: 10.1016/S1474-4422(19)30435-1 (2020).

36. Dumitrescu, L. et al. Genetic variants and functional pathways associated with resilience to alzheimer’s disease. Brain 143, 2561–2575, DOI: 10.1093/brain/awaa209 (2020).

37. Hawrylycz, M. J. et al. An anatomically comprehensive atlas of the adult human brain transcriptome. Nature 489, 391–399, DOI: 10.1038/nature11405 (2012).

38. Markello, R. D. et al. Standardizing workflows in imaging transcriptomics with the abagen toolbox. Elife 10, DOI: 10.7554/eLife.72129 (2021).

39. Freeze, B., Acosta, D., Pandya, S., Zhao, Y. & Raj, A. Regional expression of genes mediating trans-synaptic alpha-synuclein transfer predicts regional atrophy in parkinson disease. Neuroimage Clin 18, 456–466, DOI: 10.1016/j.nicl.2018.01.009 (2018).

40. Pandya, S. et al. Modeling seeding and neuroanatomic spread of pathology in amyotrophic lateral sclerosis. Neuroimage 251, 118968, DOI: 10.1016/j.neuroimage.2022.118968 (2022).

41. Arnatkeviciute, A., Fulcher, B. D. & Fornito, A. A practical guide to linking brain-wide gene expression and neuroimaging data. Neuroimage 189, 353–367, DOI: 10.1016/j.neuroimage.2019.01.011 (2019).

42. Okamura, N. et al. The development and validation of tau pet tracers: current status and future directions. Clin Transl Imaging 6, 305–316, DOI: 10.1007/s40336-018-0290-y (2018).

43. Chappell, S. et al. Observations of extensive gene expression differences in the cerebellum and potential relevance to alzheimer’s disease. BMC Res Notes 11, 646, DOI: 10.1186/s13104-018-3732-8 (2018).

44. Xia, M., Wang, J. & He, Y. Brainnet viewer: a network visualization tool for human brain connectomics. PLoS One 8, e68910, DOI: 10.1371/journal.pone.0068910 (2013).

45. Ge, S. X., Jung, D. & Yao, R. ShinyGO: a graphical gene-set enrichment tool for animals and plants. Bioinformatics 36, 2628–2629, DOI: 10.1093/bioinformatics/btz931 (2019).

46. Frost, B. & Diamond, M. I. Prion-like mechanisms in neurodegenerative diseases. Nat Rev Neurosci 11, 155–9 (2010).

47. Foster, E. M., Dangla-Valls, A., Lovestone, S., Ribe, E. M. & Buckley, N. J. Clusterin in alzheimer’s disease: Mechanisms, genetics, and lessons from other pathologies. Front Neurosci 13, 164, DOI: 10.3389/fnins.2019.00164 (2019).

48. Linden, R. The biological function of the prion protein: a cell surface scaffold of signaling modules. Front Mol Neurosci 10, 909–913, DOI: 10.3389/fnmol.2017.00077 (2017).

49. Janson, J. et al. Increased risk of type 2 diabetes in alzheimer disease. Diabetes 53, 474–81, DOI: 10.2337/diabetes.53.2.474 (2004).

50. Sobue, A. et al. Microglial gene signature reveals loss of homeostatic microglia associated with neurodegeneration of alzheimer’s disease. Acta Neuropathol Commun 9, 1, DOI: 10.1186/s40478-020-01099-x (2021).

51. Humphries, A. D. et al. Dissection of the mitochondrial import and assembly pathway for human tom40. J Biol Chem 280, 11535–43, DOI: 10.1074/jbc.M413816200 (2005).

52. Kulminski, A. M. et al. Independent associations of tomm40 and apoe variants with body mass index. Aging Cell 18, e12869, DOI: 10.1111/acel.12869 (2019).

53. Lopes, K. P. et al. Genetic analysis of the human microglial transcriptome across brain regions, aging and disease pathologies. Nat Genet. 54, 4–17, DOI: 10.1038/s41588-021-00976-y (2022).

54. Tesi, N. et al. Cognitively healthy centenarians are genetically protected against alzheimer’s disease specifically in immune and endo-lysosomal systems. medRxiv DOI: 10.1101/2023.05.16.23290049 (2023).

