## Supplemental information for "Selective vulnerability and resilience to Alzheimer’s disease tauopathy as a function of genes and the connectome"

#### Supplemental Tables

**Table S1: Genes implicated in tau selective vulnerability with function related to the brain's network.**

| Gene symbol | Gene name | $\mathbf{R}_{observed}$ | $\mathbf{R}_{residual}$ |
|---|---|---|---|
| ABCA1 | ATP binding cassette subfamily A member 1 | 0.3046 | 0.0653 |
| ANK3 | Ankyrin 3 | 0.2606 | 0.23 |
| APOE | Apolipoprotein E | 0.3507 | 0.1436 |
| BAG2 | BCL2 associated gene cochaperone | 0.1569 | 0.1159 |
| BDNF | Brain-derived neurotrophic factor | 0.0877 | 0.2814 |
| BIN1 | Bridging integrator 1 | 0.2587 | 0.0634 |
| CASP7 | Caspase 7 | 0.2766 | 0.0856 |
| CD33 | Siglec-3 | 0.0673 | 0.0153 |
| CX3CR1 | C-X3-C motif chemokine receptor 1 | 0.1008 | 0.0121 |
| DDIT4 | DNA damage inducible transcript 4 | 0.1298 | -0.0258 |
| DNAJB4 | DnaJ heat shock protein family (Hsp40) member B4 | 0.122 | 0.0527 |
| FERMT2 | Fermitin family homolog 2 | 0.2334 | 0.0435 |
| GRID2 | Glutamate ionotropic receptor delta type subunit 2 | 0.135 | 0.0323 |
| GRIN2B | Glutamate receptor ionotropic, NMDA 2B | 0.0665 | 0.1677 |
| HSP90AB1 | Heat shock protein 90 alpha family class B member 1 | 0.1698 | 0.2629 |
| IL1B | Interleukin-1beta | 0.0282 | -0.0183 |
| IL34 | Interleukin-34 | 0.1614 | 0.1926 |
| LRIG1 | Leucine-rich repeats and immunoglobulin-like domains protein 1 | 0.2785 | -0.0583 |
| MAPT | Microtubule associated protein tau | 0.2807 | 0.3184 |
| PLCG2 | Phospholipase C gamma 2 | 0.0726 | -0.0224 |
| PRKD3 | Serine/threonine-protein kinase D3 | 0.1722 | -0.0256 |
| SREBF1 | Sterol regulatory element binding transcription factor 1 | 0.1284 | -0.1029 |
| TNFRSF1A | Tumor necrosis factor receptor superfamily 1A | -0.0122 | 0.0933 |
| TSPOAP1 | TSPO associated protein 1 | 0.3511 | 0.405 |
| TYROBP | TYRO protein tyrosine kinase-binding protein | 0.0303 | -0.067 |
| ZNF184 | Zinc finger protein 184 | 0.0509 | 0.2037 |

**Table S2: Genes implicated in tau selective vulnerability with function independent of the brain's network.**

| Gene symbol | Gene name | $\mathbf{R}_{observed}$ | $\mathbf{R}_{residual}$ |
|---|---|---|---|
| CLU | Clusterin | 0.0494 | 0.1604 |
| COX7C | Cytochrome c oxidase subunit 7C | 0.0167 | 0.0547 |
| DLGAP2 | Disks large-associated protein 2 | -0.0978 | 0.2145 |
| DNAJA1 | DnaJ heat shock protein family (Hsp40) member A1 | 0.0335 | 0.2195 |
| DNAJB6 | DnaJ heat shock protein family (Hsp40) member B6 | 0.1253 | 0.3347 |
| DOC2A | Double C2 domain alpha | -0.0733 | 0.2753 |
| EED | Embryonic ectoderm development | 0.1068 | 0.2949 |
| EGR1 | Early growth response 1 | 0.0395 | 0.0842 |
| HSPH1 | Heat shock protein family H (Hsp110) member 1 | 0.1405 | 0.2288 |
| JAZF1 | Juxtaposed with another zinc-finger 1 | 0.1695 | 0.3882 |
| MAPK14 | Mitogen-activated protein kinase 14 | 0.2028 | 0.3521 |
| PILRA | Paired immunoglobin like type 2 receptor alpha | 0.0108 | -0.0171 |
| PLD3 | Phospholipase D3 | 0.0428 | 0.1877 |
| PRNP | Prion protein | 0.1614 | 0.4423 |
| RORB | Retinoid-related orphan nuclear receptor beta1 | 0.1444 | 0.1591 |
| SIRPA | Signal regulatory protein alpha | 0.2197 | 0.3795 |
| TSPAN14 | Tetraspanin-14 | 0.1437 | 0.2757 |
| ZBTB11 | Zinc finger and BTB domain containing 11 | -0.0398 | 0.1628 |

**Table S3: Genes implicated in tau selective resilience with function related to the brain's network.**

| Gene symbol | Gene name | $\mathbf{R}_{observed}$ | $\mathbf{R}_{residual}$ |
|---|---|---|---|
| ABCA7 | ATP binding cassette subfamily A member 7 | -0.0773 | -0.2207 |
| ADAM17 | ADAM metallopeptidase domain 17 | -0.0483 | -0.162 |
| ADAMTS1 | ADAM metallopeptidase with thrombospondin type 1 motif 1 | -0.1317 | -0.1406 |
| ANKH | ANKH inorganic pyrophosphate transport regulator | -0.2017 | -0.1413 |
| APP | Amyloid beta precursor protein | -0.1016 | -0.0757 |
| ATP8B1 | ATPase phospholipid transporting 8B1 | -0.3076 | -0.1775 |
| BACE1 | Beta secretase | -0.287 | -0.3143 |
| BHLHE40 | Basic helix-loop-helix family member E40 | -0.1351 | -0.0838 |
| CEBPB | CCAAT enhancer-binding protein beta | -0.1318 | -0.026 |
| CREB3L4 | CAMP responsive element binding protein 3 like 4 | -0.1529 | -0.3031 |
| FOXF1 | Forkhead box protein F1 | -0.2361 | -0.2598 |
| HS3ST2 | Heparan sulfate glucosamine 3-O-sulfotransferase 2 | -0.3434 | 0.0119 |
| HS3ST5 | Heparan sulfate glucosamine 3-O-sulfotransferase 5 | -0.4517 | -0.4323 |
| HSPA1L | Heat shock 70 kDa protein 1-like | -0.0935 | 0.1511 |
| HSPA8 | Heat shock protein family A (Hsp70) member 8 | -0.0442 | -0.0935 |
| IER5 | Immediate early response gene 5 | -0.1164 | 0.0332 |
| NCK2 | Non-catalytic region of tyrosine kinase adaptor protein 2 | -0.1814 | -0.1076 |
| NEAT1 | Nuclear enriched abundant transcript 1 | -0.1651 | -0.2933 |
| PER1 | Period circadian regulator 1 | -0.0793 | 0.0397 |
| PTK2B | Protein-tyrosine kinase 2-beta | -0.0815 | 0.1101 |
| RAB10 | Ras associated protein | -0.1326 | -0.0649 |
| SHARPIN | SHANK associated RH domain interactor | -0.1893 | -0.3144 |
| SLC44A1 | Choline transporter-like protein 1 | -0.3097 | -0.2622 |
| SOD1 | Superoxide dismutase type 1 | -0.112 | -0.0437 |
| SORL1 | Sortilin related receptor 1 | -0.1765 | -0.0249 |
| SORT1 | Sortilin | -0.3389 | -0.2089 |
| SPI1 | Spi-1 proto-oncogene | -0.0739 | -0.0689 |
| SPP1 | Secreted phosphoprotein 1 | -0.2573 | -0.3841 |
| TMEM41A | Transmembrane protein 41A | -0.0086 | -0.1012 |
| TOMM40 | Translocase of outer mitochondrial membrane 40 | -0.206 | -0.1606 |
| TREM2 | Triggering receptor expressed on nyeloid cells 2 | -0.0641 | -0.1095 |
| WDR12 | WD repeat domain 12 | -0.1563 | -0.0706 |
| WDR81 | WD repeat domain 81 | -0.2372 | -0.288 |
| ZC3H10 | Zinc finger CCCH domain-containing protein 10 | -0.1812 | -0.2083 |
| ZCWPW1 | Zinc finger CW-type and PWWP domain containing 1 | -0.1865 | -0.2296 |

**Table S4: Genes implicated in tau selective resilience with function independent of the brain's network.**

| Gene symbol | Gene name | $\mathbf{R}_{observed}$ | $\mathbf{R}_{residual}$ |
|---|---|---|---|
| APH1B | Aph-1 homolog B, gamma-secretase subunit | -0.0617 | -0.2775 |
| CD59 | Codes for membrane inhibitor of reactive lysis (MIRL) or protectin | -0.0183 | -0.1496 |
| EPDR1 | Ependymin-related protein | -0.1332 | -0.2534 |
| FBXL7 | F-box and leucine rich repeat protein 7 | -0.242 | -0.2686 |
| HBP1 | HMG-box transcription factor 1 | 0 | -0.166 |
| HSP90AA1 | Heat shock protein 90 alpha family class A member 1 | -0.0306 | -0.045 |
| IDUA | Alpha-L-iduronidase | -0.0295 | -0.1266 |
| INPP5D | Inositol polyphosphate-5-phosphatase D | -0.0219 | -0.1189 |
| MINDY2 | MINDY lysine 48 deubiquitinase 2 | -0.1373 | -0.4097 |
| NYAP1 | Neuronal tyrosine phosphorylated phosphoinositide-3-kinase adaptor 1 | -0.0168 | -0.0675 |
| PICALM | Phosphatidylinositol binding clathrin assembly protein | -0.135 | -0.2953 |
| PSEN2 | Presenilin 2 | -0.0016 | -0.0852 |
| PVR | Poliovirus receptor | -0.0223 | -0.2013 |
| REST | RE1 silencing transcription factor | -0.0757 | -0.2466 |
| SERTAD1 | SERTA domain-containing protein 1 | -0.2205 | -0.3469 |
| SNX1 | Sorting nexin 1 | -0.3904 | -0.4054 |
| SPDYE3 | Speedy/RINGO cell cycle regulator family member E3 | -0.1993 | -0.038 |
| TNIP1 | TNFAIP3 interacting protein 1 | -0.2255 | -0.3271 |
| TPCN1 | Two pore segment channel 1 | -0.0775 | -0.2404 |
| USP6NL | USP6 N-terminal like | -0.0547 | -0.2609 |
| ZBTB7A | Zinc finger and BTB domain containing 7A | -0.18 | -0.1839 |

**Table S5: Genes implicated in tau selective vulnerability genes, network-related (SV-NR).**

| Gene symbol | Gene description | $\mathbf{R}_{observed}$ | $\mathbf{R}_{residual}$ |
|---|---|---|---|
| ABCA1 | ATP binding cassette subfamily A member 1 | 0.21308 | 0.074728 |
| ANK3 | Ankyrin 3 | 0.23852 | 0.16996 |
| APOE | Apolipoprotein E | 0.24117 | 0.14028 |
| BAG2 | BCL2 associated gene cochaperone | 0.12568 | 0.065776 |
| BDNF | Brain-derived neurotrophic factor | 0.14899 | 0.13736 |
| BIN1 | Bridging integrator 1 | 0.17911 | 0.12167 |
| CASP7 | Caspase 7 | 0.18712 | 0.12282 |
| CD33 | Siglec-3 | 0.05179 | 0.005743 |
| CX3CR1 | C-X3-C motif chemokine receptor 1 | 0.072517 | 0.032729 |
| DDIT4 | DNA damage inducible transcript 4 | 0.092044 | 0.0098711 |
| DNAJB4 | DnaJ heat shock protein family (Hsp40) member B4 | 0.058489 | 0.0344 |
| FERMT2 | Fermitin family homolog 2 | 0.13915 | 0.064008 |
| GRID2 | Glutamate ionotropic receptor delta type subunit 2 | 0.066173 | 0.040078 |
| GRIN2B | Glutamate receptor ionotropic, NMDA 2B | 0.16164 | 0.10224 |
| HSP90AB1 | Heat shock protein 90 alpha family class B member 1 | 0.19464 | 0.18052 |
| IL1B | Interleukin-1beta | 0.054025 | -0.027229 |
| IL34 | Interleukin-34 | 0.19711 | 0.15682 |
| LRIG1 | Leucine-rich repeats and immunoglobulin-like domains protein 1 | 0.13553 | 0.020943 |
| MAPT' | Microtubule associated protein tau | 0.327 | 0.26111 |
| PLCG2 | Phospholipase C gamma 2 | 0.020821 | -0.029482 |
| PRKD3 | Serine/threonine-protein kinase D3 | 0.032595 | 0.0085648 |
| SREBF1 | Sterol regulatory element binding transcription factor 1 | 0.055009 | -0.023822 |
| TNFRSF1A | Tumor necrosis factor receptor superfamily 1A | 0.06333 | 0.045585 |
| TSPOAP1 | TSPO associated protein 1 | 0.36013 | 0.29975 |
| ZNF184 | Zinc finger protein 184 | 0.12686 | 0.098927 |

**Table S6: Genes implicated in tau selective vulnerability genes, network-independent (SV-NI).**

| Gene symbol | Gene description | $\mathbf{R}_{observed}$ | $\mathbf{R}_{residual}$ |
|---|---|---|---|
| CLU | Clusterin | 0.028209 | 0.10724 |
| COX7C | Cytochrome c oxidase subunit 7C | 0.012254 | 0.069804 |
| DLGAP2 | Disks large-associated protein 2 | 0.10997 | 0.1178 |
| DNAJA1 | DnaJ heat shock protein family (Hsp40) member A1 | 0.044356 | 0.11833 |
| DNAJB6 | DnaJ heat shock protein family (Hsp40) member B6 | 0.1584 | 0.25025 |
| DOC2A | Double C2 domain alpha | 0.11973 | 0.14365 |
| EED | Embryonic ectoderm development' | 0.13294 | 0.19792 |
| HSPH1 | Heat shock protein family H (Hsp110) member 1 | 0.14073 | 0.20634 |
| JAZF1 | Juxtaposed with another zinc-finger 1 | 0.21153 | 0.28957 |
| MAPK14 | Mitogen-activated protein kinase 14 | 0.25273 | 0.28112 |
| PLD3 | Phospholipase D3 | 0.058553 | 0.13481 |
| PRNP | Prion protein | 0.22599 | 0.30363 |
| RORB | Retinoid-related orphan nuclear receptor beta1 | 0.087912 | 0.13061 |
| SIRPA | Signal regulatory protein alpha | 0.25473 | 0.27699 |
| TSPAN14 | Tetraspanin-14 | 0.13576 | 0.17019 |
| ZBTB11 | Zinc finger and BTB domain containing 11 | 0.040812 | 0.12968 |

**Table S7: Genes implicated in tau selective resilience genes, network-related (SR-NR).**

| Gene symbol | Gene description | $\mathbf{R}_{observed}$ | $\mathbf{R}_{residual}$ |
|---|---|---|---|
| ADAM17 | ADAM metallopeptidase domain 17 | -0.098585 | -0.070977 |
| ADAMTS1 | ADAM metallopeptidase with thrombospondin type 1 motif 1 | -0.15742 | -0.12406 |
| ANKH | ANKH inorganic pyrophosphate transport regulator | -0.17739 | -0.097602 |
| APP | Amyloid beta precursor protein | -0.11346 | -0.095792 |
| ATP8B1 | ATPase phospholipid transporting 8B1 | -0.25988 | -0.15256 |
| BACE1 | Beta secretase | -0.23778 | -0.19703 |
| BHLHE40 | Basic helix-loop-helix family member E40 | -0.14451 | -0.061721 |
| CEBPB | CCAAT enhancer-binding protein beta | -0.16602 | -0.052235 |
| EGR1 | Early growth response 1 | -0.032345 | 0.053742 |
| FBXL7 | F-box and leucine rich repeat protein 7 | -0.21009 | -0.20971 |
| FOXF1 | Forkhead box protein F1 | -0.27287 | -0.21496 |
| HS3ST2 | Heparan sulfate glucosamine 3-O-sulfotransferase 2 | -0.18473 | -0.022809 |
| HS3ST5 | Heparan sulfate glucosamine 3-O-sulfotransferase 5 | -0.43595 | -0.30733 |
| HSPA1L | Heat shock 70 kDa protein 1-like | -0.070969 | 0.058711 |
| HSPA8 | Heat shock protein family A (Hsp70) member 8 | -0.11592 | -0.067507 |
| IER5 | Immediate early response gene 5 | -0.052307 | 0.034874 |
| NCK2 | Non-catalytic region of tyrosine kinase adaptor protein 2 | -0.15625 | -0.09195 |
| NEAT1 | Nuclear enriched abundant transcript 1 | -0.22672 | -0.19114 |
| PER1 | Period circadian regulator 1 | -0.11684 | 0.00057417 |
| PICALM | Phosphatidylinositol binding clathrin assembly protein | -0.2115 | -0.20388 |
| PILRA | Paired immunoglobin like type 2 receptor alpha | -0.043284 | -0.0074845 |
| PTK2B | Protein-tyrosine kinase 2-beta | -0.051413 | 0.017931 |
| RAB10 | Ras associated protein | -0.041125 | -0.016743 |
| SHARPIN | SHANK associated RH domain interactor | -0.2613 | -0.20668 |
| SLC44A1 | Choline transporter-like protein 1 | -0.2647 | -0.18726 |
| SOD1 | Superoxide dismutase type 1 | -0.11809 | -0.040249 |
| SORL1 | Sortilin related receptor 1 | -0.18062 | -0.02417 |
| SORT1 | Sortilin | -0.194 | -0.15617 |
| SPDYE3 | Speedy/RINGO cell cycle regulator family member E3 | -0.048785 | -0.04258 |
| SPI1 | Spi-1 proto-oncogene | -0.060603 | -0.022824 |
| SPP1 | Secreted phosphoprotein | -0.27779 | -0.24267 |
| TMEM41A | Transmembrane protein 41A | -0.10611 | -0.048649 |
| TNIP1 | TNFAIP3 interacting protein 1 | -0.21999 | -0.21588 |
| TOMM40 | Translocase of outer mitochondrial membrane 40 | -0.206 | -0.1375 |
| TREM2 | Triggering receptor expressed on nyeloid cells 2 | -0.08159 | -0.077842 |
| WDR12 | WD repeat domain 12 | -0.16929 | -0.07098 |
| WDR81 | WD repeat domain 81 | -0.23926 | -0.18982 |
| ZBTB7A | Zinc finger and BTB domain containing 7A | -0.16307 | -0.15835 |
| ZC3H10 | Zinc finger CCCH domain-containing protein 10 | -0.1797 | -0.13663 |
| ZCWPW1 | Zinc finger CW-type and PWWP domain containing 1 | -0.15403 | -0.11946 |

**Table S8: Genes implicated in tau selective resilience genes, network-independent (SR-NI).**

| Gene symbol | Gene description | $\mathbf{R}_{observed}$ | $\mathbf{R}_{residual}$ |
|---|---|---|---|
| ABCA7 | ATP binding cassette subfamily A member 7 | -0.15373 | -0.16045 |
| APH1B | Aph-1 homolog B, gamma-secretase subunit | -0.12926 | -0.19637 |
| CD59 | Codes for membrane inhibitor of reactive lysis (MIRL) or protection | -0.048941 | -0.078919 |
| CREB3L4 | CAMP responsive element binding protein 3 like 4 | -0.17348 | -0.17794 |
| EPDR1 | Ependymin-related protein | -0.10572 | -0.17486 |
| HBP1 | HMG-box transcription factor 1 | -0.07208 | -0.093695 |
| HSP90AA1 | Heat shock protein 90 alpha family class A member 1 | -0.0087477 | -0.052352 |
| IDUA | Alpha-L-iduronidase | -0.072854 | -0.080833 |
| INPP5D | Inositol polyphosphate-5-phosphatase D | -0.057621 | -0.087045 |
| MINDY2 | MINDY lysine 48 deubiquitinase 2 | -0.1986 | -0.2583 |
| NYAP1 | Neuronal tyrosine phosphorylated phosphoinositide-3-kinase adaptor 1 | -0.03119 | -0.081209 |
| PSEN2 | Presenilin 2 | -0.024433 | -0.047915 |
| PVR | Poliovirus receptor | -0.10664 | -0.14005 |
| REST | RE1 silencing transcription factor | -0.078416 | -0.15484 |
| SERTAD1 | SERTA domain-containing protein 1 | -0.18314 | -0.22359 |
| SNX1 | Sorting nexin 1 | -0.32055 | -0.32801 |
| TPCN1 | Two pore segment channel 1 | -0.047302 | -0.14988 |
| TYROBP | TYRO protein tyrosine kinase-binding protein | -0.0033731 | -0.040144 |
| USP6NL | USP6 N-terminal like | -0.11947 | -0.18662 |

### Supplemental Figures

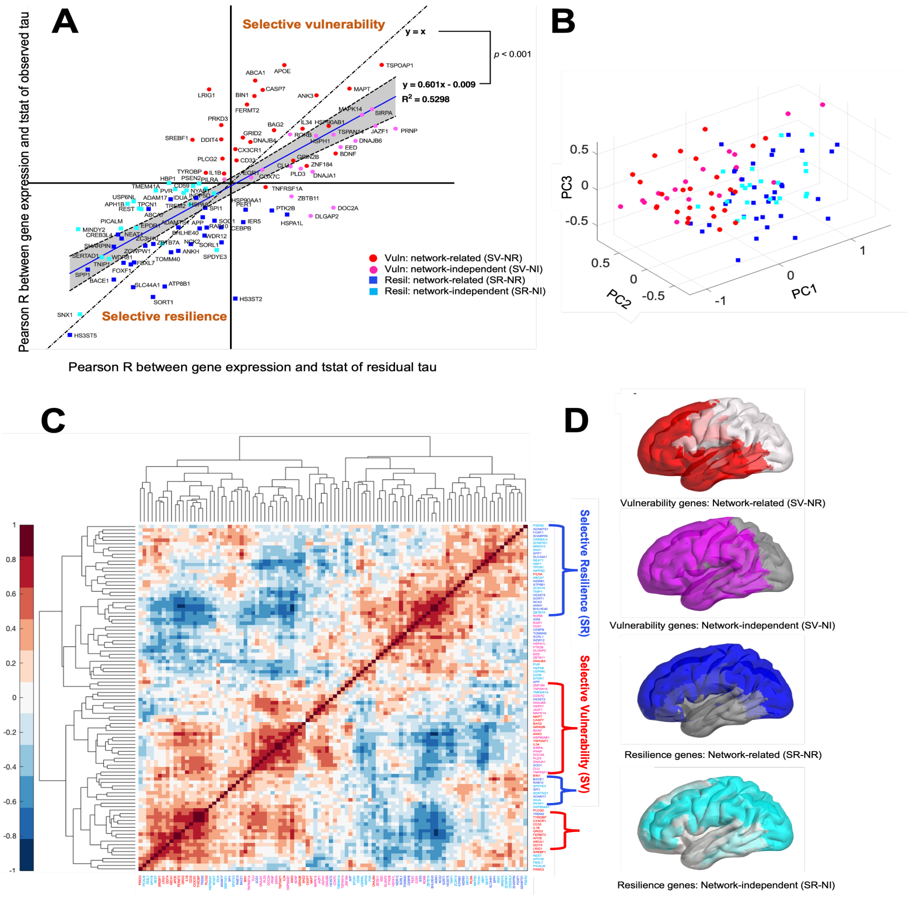


**Figure S1: Group level analysis of genes implicated in Alzheimer's disease using t-scores.**

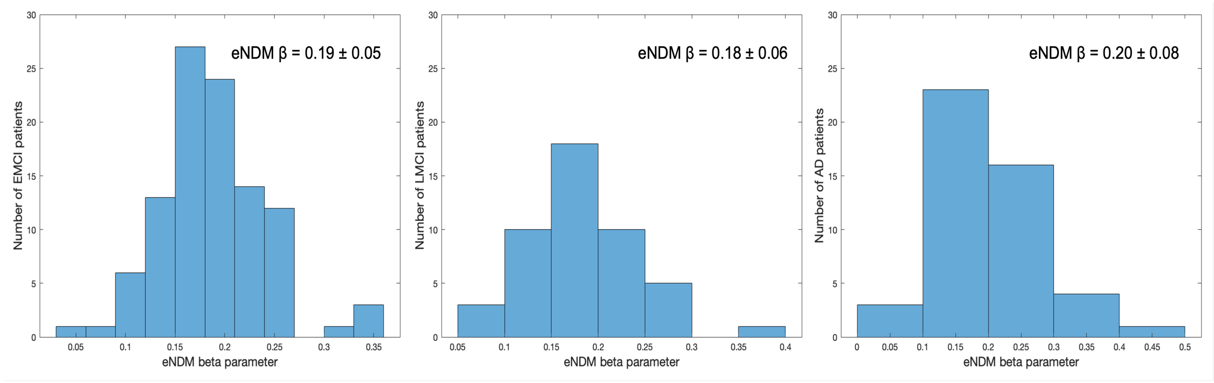


**Figure S2: Distribution of the eNDM diffusion parameter, β, across patients.** Histograms depict the distribution (mean ± 1SD) of the eNDM diffusion parameter, β, in the EMCI, LMCI, and AD groups. The values are in agreement across the three groups.
